# Far-reaching displacement effects of artificial light at night in a North American bat community

**DOI:** 10.1101/2023.10.11.561893

**Authors:** Chad L. Seewagen, Julia Nadeau-Gneckow, Amanda M. Adams

**Affiliations:** Great Hollow Nature Preserve & Ecological Research Center, New Fairfield, Connecticut, USA; Bat Conservation International, Austin, Texas, USA; Department of Biology, Texas A&M University, College Station, Texas, USA

**Keywords:** artificial light at night, light pollution, LED, habitat loss, bat community, *Myotis lucifugus*

## Abstract

Artificial light at night (ALAN) is a global pollutant that disrupts circadian rhythmicity and a broad range of physiological processes and behaviors in animals. However, ALAN sensitivity can vary greatly even among closely related species and urgently needs study for much of the world’s nocturnal wildlife, including bats. While an increasing number of bat species have been assessed for light tolerance in recent years, the spatial extent of ALAN’s influence on bats has received little attention. This information need is a barrier to the protection of bats from ALAN in land-use planning and policy, and the development of best practices that effectively buffer bat habitat from light trespass. To help address this information gap for North America, we experimentally tested the distances up to which ALAN affects presence and activity of light-averse little brown bats (*Myotis lucifugus*) and big brown bats (*Eptesicus fuscus*), and the composition of a foraging bat assemblage in Connecticut, USA. We used three residential-scale, white, LED floodlights to expose bat foraging habitat to ALAN and compared acoustic activity of bats at distances of 0, 25, 50, and 75 m from the lights between nights when the lights were on versus off. Little brown bats were present on significantly fewer light than dark nights at every distance. Lighting significantly reduced little brown bat activity overall and at the farthest location from the lights (75 m), where it was only 43% of dark-night activity despite 0 lx of illuminance. Presence of big brown bats was not significantly affected at any distance. Big brown bat activity on light nights averaged 48-75% of dark-night activity at each distance but was significantly lower only at 0 m. Community composition on dark and light nights had an average dissimilarity of 38% and significantly differed at 0 m and 25 m. We conclude the type of ALAN used in our study has a disturbance radius of at least 75 m for the little brown bat and up to 25 m for the big brown bat, with a resulting influence on community composition for up to 50 m. Cumulative habitat loss for the imperiled little brown bat caused by ALAN could therefore be substantial. We urge planners and natural resources regulators to consider these footprints when evaluating indirect impacts to bat habitat from current and future sources of ALAN across these species’ ranges.

## 1. INTRODUCTION

Artificial light at night (ALAN) is a global pollutant that affects nearly a quarter of the world’s land surface and is spreading rapidly in both industrialized and developing nations in concert with human population growth, urbanization, and advancements in lighting technology and cost efficiency (Holker et al. 2010a; Falchi et al. 2016; Kyba et al. 2017a, 2023).

Measurements of sky glow indicate outdoor light emissions have risen worldwide by a yearly average of almost 10% in just the past decade (Kyba et al. 2023), with a growing contribution from broad-spectrum lights, such as LED, that may be worsening ecological impacts (Pawson and Bader 2014, Sánchez de Miguel et al. 2022). With these trends expected to continue, there is mounting concern for ALAN’s impacts on biodiversity and increasing evidence for adverse effects on a wide range of animal taxa (Holker et al. 2010b, Knop et al. 2017, Falcon et al. 2020, Burt et al. 2023). ALAN’s disruption of natural day-night cycles can alter an animal’s physiology and cause maladaptive changes in behavior that ultimately reduce fitness and survival (reviewed by Gaston et al. 2013, Falcon et al. 2020). By affecting some species and not others, ALAN can also drive shifts in species interactions, competitive balance, and the composition of animal communities, potentially leading to ecosystem-level consequences (Davies et al. 2013, Manfrin et al. 2017, Falcon et al. 2020). A better understanding of how organisms respond to ALAN has thus become a conservation priority and pressing need in the search for effective and practical measures to minimize its ecological impacts (Holker et al. 2010b, Gaston et al. 2014, Schroer et al. 2020).

The degradation of nightscapes by ALAN is expected to have the greatest effects on nocturnal animals, such as bats, whose unique sensory and behavioral adaptations to darkness partition them from the niches of diurnal species (Beier 2006, Rowse et al. 2016, Sanders et al. 2021). ALAN can delay bats’ emergence times, impede movement and habitat connectivity, reduce foraging activity, and alter trophic interactions (e.g., Boldough et al. 2007; Hale et al. 2015; Cravens and Boyles 2018, 2019; Azam et al. 2018, Russo et al. 2019). The effects of ALAN on bats are often highly species-specific, however, with some species avoidant of lights and others attracted to them by associated concentrations of insect prey (Stone et al. 2015, Rowse et al. 2016). This attraction-repulsion dynamic and the barrier ALAN poses to some species’ space-use but not others can be strong enough to alter bat community composition at local to landscape scales (Azam et al. 2016, Schoeman 2016, Seewagen and Adams 2021).

Most bats that are currently known to be light-averse belong to the genus *Myotis* (Stone et al. 2009, Rowse et al. 2016). Widespread light avoidance among *Myotis* bats is thought to be due to their relatively slow flight speeds, which may increase perceived predation risk in lit environments more so than for faster-flying, light-tolerant species (Jones and Rydell 1994, Stone et al. 2015). In North America, *Myotis* species and some other cave-hibernating bats are experiencing extreme population declines of more than 90% due to White-nose Syndrome (WNS; Cheng et al. 2021). Minimizing anthropogenic stressors to these species has therefore become of heightened importance to their conservation until practical ways to alleviate the effect of WNS can be identified or there is sufficient selection for disease resistance before populations are too bottlenecked to recover (USFWS 2015, McClure et al. 2022).

Many *Myotis* species require high habitat connectivity for commuting among foraging and roosting areas (Henderson and Broders 2008, Browning et al. 2021, Cable et al. 2021, Frantz et al. 2022). Therefore, ALAN is likely to be an agent of fragmentation and habitat loss for light-averse bats (Ditmer et al. 2021). Yet, the footprint of disturbance to bat habitat caused by ALAN is largely unknown because the distances up to which ALAN displaces light-averse bats have not been studied for most species. Some European bats have been shown to avoid lights 8-50 m from the source (Azam et al. 2018, Barre et al. 2021a,b); otherwise, to our knowledge, the size of the effect zone surrounding lights remains unexplored for the world’s more than 1400 other bat species. This knowledge gap is an impediment to the protection of bats in land-use planning and policy (Pauwels et al. 2019, Schroer et al. 2020). To help address this information need for North America, we built upon previous research where we documented significant LED light avoidance by the little brown bat (*Myotis lucifugus*) and big brown bat (*Eptesicus fuscus*) at an exurban study site in Connecticut, USA (Seewagen and Adams 2021). Here, we experimentally tested the distances up to which that same LED lighting affects these species’ presence and foraging activity and the composition of the foraging bat community to understand the reach of ALAN’s influence on light-averse North American bats.

## 2. MATERIALS AND METHODS

### 2.1 Study Site

We conducted our experiment in a wetland at Great Hollow Nature Preserve in New Fairfield, Connecticut, USA (41.502320, - 73.531300). The preserve is approximately 335 ha of mostly second-growth, mature forest and is contiguous or semi-contiguous with approximately 1,330 ha of adjacent protected lands in Connecticut and New York State. Development surrounding the preserve is low-density and residential and the two roads bordering the preserve lack street lighting. As such, there are no chronic sources of ALAN to which our study site is exposed. The wetland is approximately 4.8 ha of open and emergent wetland that is partially maintained by periodic impoundment of Quaker Brook by American beavers (*Castor canadensis*). It is bordered on its eastern and western sides by mature hardwood forest, and on its northern and southern sides by a matrix of mowed lawn, wet meadow, old field, and shrubland.

### 2.2 Lighting Treatment and Recording Methods

Our experimental design was as described in Seewagen and Adams (2021) except that we deployed three additional acoustic bat recorders at 25-m intervals away from the light array. We also mounted the lights and microphones 0.5 m higher to account for an increase in vegetation height in the wetland since the previous study and we did not include a treatment in which the lighting infrastructure was removed because we previously found no effect of the infrastructure alone on bat activity. The light array consisted of three 55-watt, 4,400-lumen, white, LED utility lights (Keystone LED Lighting, Erie, CO, USA) mounted on 5-m poles linearly spaced 10 m apart from each other along the eastern edge of the wetland, facing west. The lights were angled 45° downwards and had a beam width of 120°. The lights’ spectral pattern was bimodal, with peaks at 450 nm and 590 nm (data provided by manufacturer). We powered the lights via a hardwire connection to the preserve’s administrative building, approximately 115 m away. The lighting treatment was approved by the University of Connecticut’s Institutional Animal Care and Use Committee (IACUC Protocol # E20-004).

Adjacent to the center light pole (hereafter 0 m) and at distances of 25, 50, and 75 m north of the northernmost light pole, we deployed Song Meter SM4BAT FS ultrasonic recorders with SMM-UI directional microphones (Wildlife Acoustics Inc., Maynard, MA, USA) from 27 June to 11 August, 2022. The microphones were mounted on metal poles 4 m above ground, along the wetland’s eastern edge, pointing west and angled approximately 45° towards the open sky. We configured the recorders to collect 3-s, full-spectrum, triggered sound files at a sampling rate of 384 kHz and gain of 12 dB, for 3 h each night, beginning at sunset. We randomly assigned each night to a light or dark treatment (lights on, lights off) by flipping a coin (*N* = 18- 21 recorder-nights per treatment, per location after omitting nights with precipitation and equipment failure). Ground-level horizontal illuminance generated by the lights was 24 lx at the 0 m recording location, 2 lx at 25 m, 1 lx at 50 m, and 0 lx at 75 m (LX-1010B photometer, Tondaj Instruments, Shenzhen, China).

We processed recordings using the Bats of Connecticut classifier in Kaleidoscope Pro version 5.4.8, set to neutral (Wildlife Acoustics Inc., Maynard, MA, USA). We accepted auto-classified species identifications when the maximum likelihood probability value was ≤ 0.05. We then manually assessed the recordings from all automated identifications of *Myotis* species to either confirm identity, reclassify them as a different species, or assign them as an unknown *Myotis* or other unknown species. We discarded recordings that could only be classified as an unknown bat species and assumed all unknown *Myotis* recordings to be those of the little brown bat because the other *Myotis* species of the region (*M. sodalis, M. septentrionalis, M. lebeii*) are either unknown to occur or extremely rare in our study area (Hammerson 2004, Seewagen and Adams 2021, CTDEEP 2022).

### 2.3 Statistical Analyses

We conducted all statistical analyses in RStudio 2022.07.1 except for community composition analyses, which we conducted in PAST 3.13 (Hammer and Harper 2001). We used an α level of 0.05 for all tests. Four species were detected with enough regularity to examine treatment and distance effects on their presence and activity: little brown bat, big brown bat, eastern red bat (*Lasiurus borealis*), and hoary bat (*Lasiurus cinereus*). The only other species we identified with significant confidence during the experiment was the silver-haired bat (*Lasionycteris noctivagans*), which occurred only sporadically across both treatments.

We used two-tailed Fisher’s exact tests to compare the number of light and dark nights a species was present (≥ 1 file recorded) at a given location. We measured bat activity by converting the number of sound files of each species, on each night, at each location to an activity index that represented a count of 1-min segments in the recording period during which the species was detected (Miller 2001). We then tested the effects of lighting treatment, recording location, and their interaction on bat activity using quasi-Poisson (log link) generalized linear models (GLM) for big brown, eastern red, and hoary bat to account for overdispersion (Zeileis et al. 2008, Beckerman et al. 2017), and a zero-inflated negative binomial (ZINB) model (*pcsl* package; Zeileis et al. 2008) for little brown bat to account for overdispersion and an abundance of zeros (Zuur et al. 2009, Hilbe 2012). We detected no little brown bats at the 0-m location when the lights were on (i.e., there were no non-zero values), so we could run that species’ model on activity data from only the 25-, 50-, and 75-m recording locations. We did not account for temperature or wind speed in our models because they did not differ between treatments (temperature: *t*51 = 0.72, *P* = 0.48; wind: χ^2^ = 0.10, *P* = 0.76).

We tested whether there was a treatment effect on bat activity that depended on distance from the light source and whether bat activity differed overall among recording locations by comparing full models to nested models from which the interaction term and recording location were sequentially removed. We compared full and nested models to each other using *F*-tests for the quasi-Poisson GLMs and likelihood ratio tests (*lmtest* package) for the ZINB model and considered a missing term unimportant for explaining variation in bat activity when the comparison was nonsignificant (Zeileis et al. 2008, Zuur et al. 2009). We then used the *emmeans* package (Lenth et al., 2023) to make pairwise, planned contrasts of bat activity between treatments at each recording distance.

We used two-way permutational multivariate analysis of variance (PERMANOVA) with Bray-Curtis distances to test the effects of lighting treatment, recording location, and their interaction on species composition (Anderson 2001, 2017). We then used one-way PERMANOVA to make pairwise, planned contrasts of bat species composition between treatments at each recording distance. Lastly, we used analysis of similarity percentage (SIMPER) to measure each species’ relative contribution to any observed differences in species composition.

## 3. RESULTS

### 3.1 Presence/Absence

In the logistic component of the ZINB model, treatment did not significantly affect the probability of detecting zero little brown bats (*F* = -1.30, *P* = 0.19), after dropping recording distance (*F* = 1.68, *P* = 0.43) and the interaction (*F* = 2.04, *P* = 0.36). However, little brown bats were present on significantly fewer light than dark nights at every recording location (Fisher’s exact tests, all *P* ≤ 0.05) (Table 1). When the lights were on, no little brown bats were detected at the 0-m location and they were detected only twice (i.e., 2 files recorded) at the 25-meter location. The presence of little brown bats on light nights increased with distance, but at 75 m, was still less than half that of dark nights (Table 1). Big brown bats, in contrast, were present at each recording distance almost every night under both treatments, with no significant effect of treatment on presence at any distance (all *P* > 0.43) (Table 1). Eastern red bat (all *P* = 1.00), hoary bat (all *P* > 0.48), and silver-haired bat (all *P* > 0.66) presence was also unaffected by treatment at all recording distances.

**Table 1.** Percentage of naturally dark (N = 18-21) and artificially lit (N = 19-20) nights bats were present at increasing distances from the light source.

|  | 0 m |  | 25 m |  | 50 m |  | 75 m |  |
| --- | --- | --- | --- | --- | --- | --- | --- | --- |
| Species | Dark | Light | Dark | Light | Dark | Light | Dark | Light |
| *Eptesicus fuscus* | 95 | 90 | 95 | 90 | 95 | 90 | 94 | 90 |
| *Lasiurus borealis* | 100 | 100 | 100 | 100 | 100 | 100 | 100 | 100 |
| *Lasiurus cinereus* | 100 | 95 | 91 | 95 | 91 | 100 | 94 | 100 |
| *Lasionycteris noctivagans* | 11 | 15 | 10 | 15 | 38 | 32 | 17 | 21 |
| *Myotis lucifugus* | 47 | 0 | 67 | 10 | 71 | 32 | 67 | 32 |

### 3.2 Foraging Activity

The activity of little brown bats on light nights averaged only 0-43% of dark-night activity at the four recording distances. Lighting significantly reduced little brown bat activity (*z* = 5.49, *P* < 0.001) regardless of distance (treatment by distance interaction: χ^2^ = 1.909, *P* = 0.385), while there was no difference in little brown bat activity among recording distances overall (i.e., both treatments combined; χ^2^ = 1.944, *P* = 0.378). In planned contrasts, little brown bats were significantly less active on light than dark nights at 25 m (*z* = 4.072, *P* = 0.007) and 75 m (*z* = 2.989, *P* = 0.033), but not at 50 m (*z* = 2.374, *P* = 0.165) (Fig. 1). Big brown bat activity on light nights averaged 48-75% of dark-night activity at the four recording distances. There was a significant negative effect of the lighting on big brown bat activity (*F*1 = 20.00, *P* < 0.001), independent of recording distance (treatment by distance interaction: *F* = 0.66, *P* = 0.579). In planned contrasts, however, big brown bats were significantly less active on light nights only at 0 m (0 m: *z* = 3.18, *P* = 0.032; 25 m: *z* = 2.35, *P* = 0.268; 50 m: *z* = 1.948, *P* = 0.518; 75 m: *z* = 1.199, *P* = 0.932; Fig. 1). Treatment had no significant effect on the activity of eastern red bats (*F*1 = 0.69, *P* = 0.41) or hoary bats (*F*1 = 2.15, *P* = 0.15).

**Figure 1.**
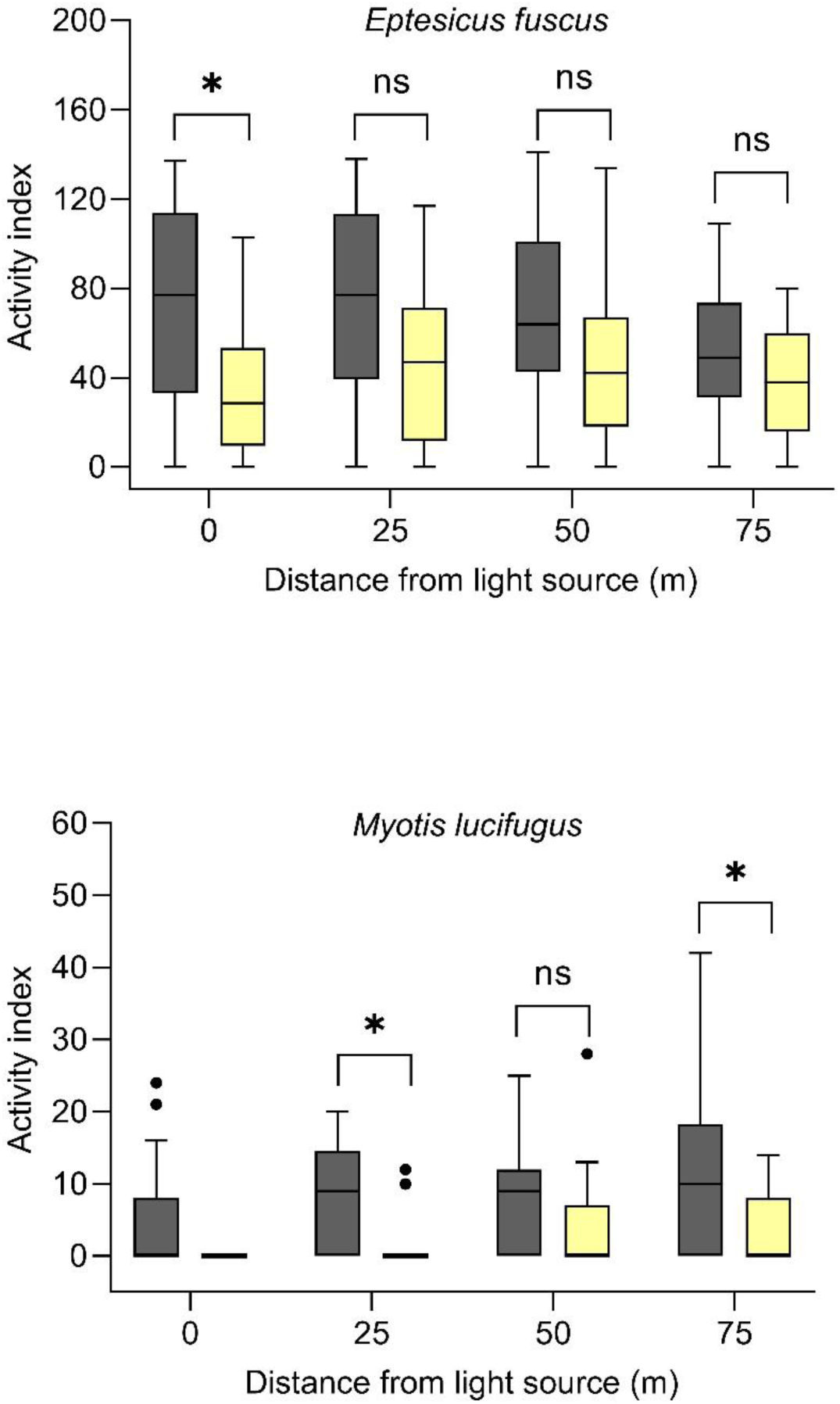
Bat activity under naturally dark (gray boxes) and artificially lit (yellow boxes) conditions, 0-75 m from the light source. Boxes show the median and 25th to 75th percentiles, whiskers represent minimum and maximum values, and black circles are outliers. Brackets show significance of planned contrasts (*P < 0.05, ns = P > 0.05, no bracket = untested because of no non-zero values in the light treatment).

### 3.3 Community Composition

Community composition was significantly different between treatments (*F* = 11.31, *P* < 0.001), independent of recording distance (treatment by distance interaction: *F* = -1.05, *P* = 0.95), and did not differ among recording distances overall (*F* = 1.55, *P* = 0.13). Reduced activity of little brown bats and big brown bats on light nights contributed 17% and 29%, respectively, to an average community dissimilarity of 38% between treatments. In planned contrasts, the lighting effect on species composition was significant at 0 m (*F* = 3.97, *P* = 0.014) and 25 m (*F* = 3.62, *P* = 0.023), approached significance at 50 m (*F* = 2.36, *P* = 0.086), and was nonsignificant at 75 m (*F* = 1.20, *P* = 0.340).

## 4. DISCUSSION

As awareness of the threat of ALAN to bats has grown rapidly in recent years and an increasing number of species have been found to be light-averse, the spatial extent of ALAN’s effects on bats has remained largely uninvestigated, particularly outside of Europe. We found residential-scale, white LED lights to have far-reaching displacement effects on two WNS-impacted North American bat species, driving shifts in overall community composition that extend well beyond the primary area of illumination. The lights had a disturbance radius of at least 75 m for the little brown bat and up to 25 m for the big brown bat, with a resulting influence on community composition up to 50 m away despite minimal to no illuminance at these distances. Our results suggest indirect habitat loss for little brown bats and big brown bats caused by the encroachment of ALAN into dark landscapes could be cumulatively substantial, with possible cascading effects on species interactions and competitive balance among the ecological communities to which they belong.

Similar effect distances as ours were found by Azam et al. (2018) among *Myotis* and *Eptesicus* species in France, where high-pressure sodium (HPS) streetlights reduced foraging activity at respective distances of 50 and 25 m. In the Netherlands, *Myotis/Plecotus* and *Eptesicus/Nyctalus* species groups were observed to alter their flight paths towards dark refugia once they were within approximately 15 m of broad-spectrum, white streetlights (Barre et al. 2021a). Pond bats (*M. dasycneme*) in the Netherlands also veered away from halogen light at approximately 15 m (Kuijper et al. 2008). Barre et al. (2021b) found *Pipistrellus* species to distance themselves an average of 10.7-15.7 m from artificially lit bridges while commonly flying over or under unlit bridges on the same urban waterway in France. From these studies and ours, it is evident that multiple types of lighting in various settings can have displacement effects that extend well beyond the light source for many bat species.

Effect distances of ALAN depend on the degree of luminance attenuation across space, which is a product of lighting technology, color, intensity, orientation, shielding, height, and other characteristics (Pauwels et al. 2019). Little brown bats and big brown bats in our study exhibited light avoidance where luminance levels were only 0-2 lx, showing that displacement effects on these species can extend into areas of darkness or near-darkness, beyond a light’s halo. Azam et al. (2018) also found avoidance of HPS streetlights by *Myotis* and *Eptesicus* species at distances where luminance had attenuated below 1 lx. Similarly, *Myotis/Plecotus* and *Eptesicus/Nyctalus* species groups in the Netherlands showed altered flight paths away from white LED streetlights at distances with corresponding light levels of 1 lx (Barre et al. 2021b). Given that moonlight varies by extremes of only ∼ 0.3 lx between full and new moons (Kyba et al. 2017b) and can influence the activity of some bat species (Saldaña-Vázquez and Munguía-Rosas 2013), it is perhaps unsurprising that even dim artificial light can affect light-averse bats. Our results suggest white LED lighting should be shielded or distanced from foraging habitats and movement corridors enough to limit trespass to 0 lx for little brown bats and less than 1 lx for big brown bats.

In non-urban landscapes of North America, where bat habitat is most abundant, ALAN from roads, energy infrastructure, public recreation areas, businesses, homes, and other sources is pervasive and expanding (Benfield et al. 2018, Boslett et al. 2021, LaSorte et al. 2022), and an increasing threat to light-averse bats. For instance, 50% of the land area of the United States is now exposed to ALAN (Falchi et al. 2016) while only 3% of its land area is urban (U.S. Census Bureau 2012). Our observed sensitivities of little brown bats and big brown bats to dim light, far from the source, have the potential to impede these species’ movements and fragment, degrade, or effectively eliminate a cumulatively significant amount of otherwise suitable habitat remaining in rural and suburban areas of their geographic ranges. The presence of big brown bats and to a lesser extent, little brown bats, in cities (Coleman and Barclay 2011, Schimpp et al. 2018, Deeley et al. 2021) suggests there could be selection for light tolerance in urban bat populations or some capacity in these species for individual or generational habituation to ALAN that would allow bats in relatively dark landscapes to gradually adapt to its encroachment (Russo et al. 2019, Seewagen and Adams 2021, Hooton et al. 2022). However, such selection or habituation might act only at a broad spatial scale while at finer scales, light-averse bats are limited to areas within cities where ALAN levels are lowest (Hale et al. 2015; Parkins et al. 2016; Barre et al. 2021a,b; 2023), leaving habitat availability extremely low relative to the total land area of a city. Interpopulation variation in light tolerance and the ability of bats to habituate to light pollution need study and should be research priorities to better understand the threats posed by ALAN to bats across the globe.

We observed no positive or negative effect of our lighting treatment on the activity of eastern red bats or hoary bats at any distance. These species appear to be indifferent to the type of lighting we studied and are perhaps unaffected by similar lighting in other contexts. Previous studies have documented an attraction of migratory tree bats, including the eastern red bat and hoary bat, to ALAN (e.g., Furlonger et al. 1987, Hickey et al. 1996), which is of concern because these species suffer the highest mortality rates at wind energy facilities (Guest et al. 2022). However, observations of eastern red and hoary bats being attracted to ALAN have mostly involved light colors (e.g., white) and technologies (e.g., mercury vapor) that tend to be more attractive to insects and light-tolerant bats than the colors (e.g., red) and modern technologies (e.g., LED) typically used for obstruction lighting on wind turbines (Bennett et al. 2014, van Grunsven et al. 2014, Lewanzik and Voigt 2017, Spoelstra et al. 2017, Guest et al. 2022, but see Voigt et al. 2018). There is currently no clear indication that lighting increases bat mortality in wind energy areas, and the attraction of tree bats to wind turbines could instead be due to bats misidentifying turbines as trees in which to roost, forage, or engage in social interactions (Guest et al. 2022). The role of ALAN in tree bat mortality at wind energy facilities warrants further investigation, however, and continues to be an active area of research.

The results of our experiment provide an important step towards understanding the spatial footprint of disturbance from ALAN to light-averse bats in non-urban landscapes that are under increasing threat of light pollution. Threatened and endangered species that are closely related to and sympatric with the little brown bat (e.g., *M. sodalis, M. septentrionalis*, *M. grisescens*) could be similarly affected by ALAN, raising the possibility that ALAN is an anthropogenic stressor that has contributed to their declines. However, much remains to be learned about how light affects North America’s bat species at individual, population, and community levels. In much the same way the effects of ALAN on bats have been progressively studied in Europe for more than two decades, research is urgently needed to assess light tolerance in North American species across a range of lighting characteristics (e.g., intensity, spectral composition) that can influence impact severity. Concurrently, we need to grow the body of evidence supporting conservation actions that strive to limit the effect of ALAN on bats. Of 15 bat conservation actions related to ALAN identified by Berthinussen et al. (2021), 10 currently have limited to no evidence of effectiveness. Better information on ALAN tolerance and the effectiveness of impact minimization measures could then be combined with geospatial light pollution and land-cover data to map and quantify the amount of habitat likely impacted by ALAN now and under future development scenarios, providing regulators and practitioners with a powerful tool to identify and protect dark spaces for bats.

## ACKNOWLEDGEMENTS

We thank Christen Long for manually reviewing and classifying bat recordings, Devaugh Fraser for lending field equipment and providing thoughtful advice on experimental design, and Wales Carter and Tina Cheng for their helpful suggestions for data analyses. Funding for the collection, analysis, and interpretation of the data for this research was provided by the Connecticut Department of Energy and Environmental Protection (personal service contract # DEPA00002070129) and Great Hollow Nature Preserve and Ecological Research Center.

